# Reconstructing the Mitochondrial Proton Motive Force Using Physics-Informed Neural Networks and Surrogate Bioenergetic Signals

**DOI:** 10.1101/2025.09.19.677356

**Authors:** Mark I.R. Petalcorin

## Abstract

The mitochondrial proton motive force (PMF) underlies ATP synthesis, metabolite transport, and energy coupling. Yet, direct measurement of PMF remains technically challenging due to probe invasiveness, calibration drift, and compartmental averaging. Here, we introduce a **physics-informed neural network (PINN)** framework that reconstructs PMF from surrogate signals including NADH, oxygen, electron flux, proton leak, and reactive oxygen species (ROS). Using a synthetic curriculum dataset derived from biophysical ranges reported in the literature, our model achieved high predictive accuracy (R^2^ ≈ 0.99, RMSE < 1 mV) under normoxia and hypoxia. SHAP-based interpretability revealed distinct feature contributions: flux and NADH dominated under normoxia, while oxygen and ROS became more influential under hypoxia. Extended analyses demonstrated that PINNs generalize robustly across cross-validation folds, preserve biophysical constraints, and can be adapted to **time-series dynamics**, capturing PMF decline and recovery during simulated hypoxia-reoxygenation. To our knowledge, this is the first application of PINNs to mitochondrial bioenergetics, bridging machine learning with the chemiosmotic theory. This proof-of-concept establishes a foundation for non-invasive PMF estimation and opens avenues for studying mitochondrial adaptation in physiology and disease.

## Introduction

The **proton motive force (PMF)** is the electrochemical gradient across the mitochondrial inner membrane that powers ATP synthesis and maintains cellular energy homeostasis (Nicholls & Ferguson, 2013). It is generated by the activity of the **electron transport chain (ETC)**, which couples substrate oxidation to proton pumping from the matrix to the intermembrane space. PMF is composed of two interdependent components: the electrical membrane potential (Δψ), generated by charge separation, and the chemical proton gradient (ΔpH), reflecting proton concentration differences (Mitchell, 1961; Zorova et al., 2018).Together, these forces drive **oxidative phosphorylation**, regulate metabolite transport, and influence **reactive oxygen species (ROS) production**, positioning PMF as a master regulator of mitochondrial and cellular physiology (Brand, 2016; Wallace, 2005).

Despite its central role, **accurate quantification of PMF remains challenging**. Experimental approaches commonly rely on **potentiometric dyes** (e.g., tetramethylrhodamine methyl ester [TMRM], JC-1) or **genetically encoded voltage indicators**. While widely used, these methods suffer from calibration drift, phototoxicity, dye aggregation, and limited temporal and spatial resolution (Brand & Nicholls, 2011; Murphy, 2009; Loew, 2015). Moreover, they typically report average signals across heterogeneous mitochondrial populations, masking **cell-to-cell and organelle-to-organelle variability** that has been shown to shape bioenergetic adaptation (Glancy et al., 2017; Lapuente-Brun et al., 2013). The technical barriers to direct PMF measurement constrain our ability to investigate its dynamics in health and disease, including conditions such as **ischemia-reperfusion injury, neurodegeneration**, and **cancer metabolism** (Chouchani et al., 2014; Zorov, Juhaszova, & Sollott, 2014).

In parallel, **machine learning (ML)** has emerged as a powerful tool to infer latent physiological states from surrogate measurements. ML approaches have been successfully applied to cardiac electrophysiology, neuroimaging, and metabolic flux prediction (Dale et al., 2010). However, standard **“black box” neural networks** pose limitations in biophysical applications: they may generate predictions that violate physical laws, and their lack of interpretability hinders mechanistic insight (Rudin, 2019). These drawbacks have spurred interest in **physics-informed neural networks (PINNs)**, which integrate mechanistic constraints into the learning process by embedding governing equations into the loss function (Raissi, Perdikaris, & Karniadakis, 2019). By enforcing consistency with established principles, PINNs achieve both predictive accuracy and mechanistic plausibility.

PINNs have been applied in diverse domains, including **fluid mechanics** (Raissi et al., 2020), **protein structure prediction** (Zhao et al., 2025), and **cardiac electrophysiology** (Sahli et al., 2020). Yet, to our knowledge, they have not been applied to **mitochondrial bioenergetics**, despite the field’s longstanding need for methods that reconcile predictive modeling with physical constraints.

Here, we present a **PINN framework for reconstructing PMF** from surrogate mitochondrial signals, including NADH concentration, oxygen availability, electron flux, proton leak, and ROS. To capture the interplay between mitochondrial bioenergetics and machine learning, we designed a framework that reconstructs the proton motive force (PMF) as the combined output of membrane potential (Δψ) and proton gradient (ΔpH) generated by complexes I-IV of the electron transport chain (Mitchell, 1961; Nicholls & Ferguson, 2013) (Figure 1A). Surrogate features including NADH, oxygen, flux, proton leak, and ROS were used as inputs to a feedforward or physics-informed neural network (PINN), which predicted PMF while respecting biophysical constraints (Raissi et al., 2019). The overall workflow (Figure 1B) spanned synthetic data generation, model training, interpretability with SHAP (Lundberg & Lee, 2017), and dynamic extensions using recurrent PINNs, thereby providing a systematic pipeline for studying mitochondrial energetics under normoxia and hypoxia.

**Figure 1.**
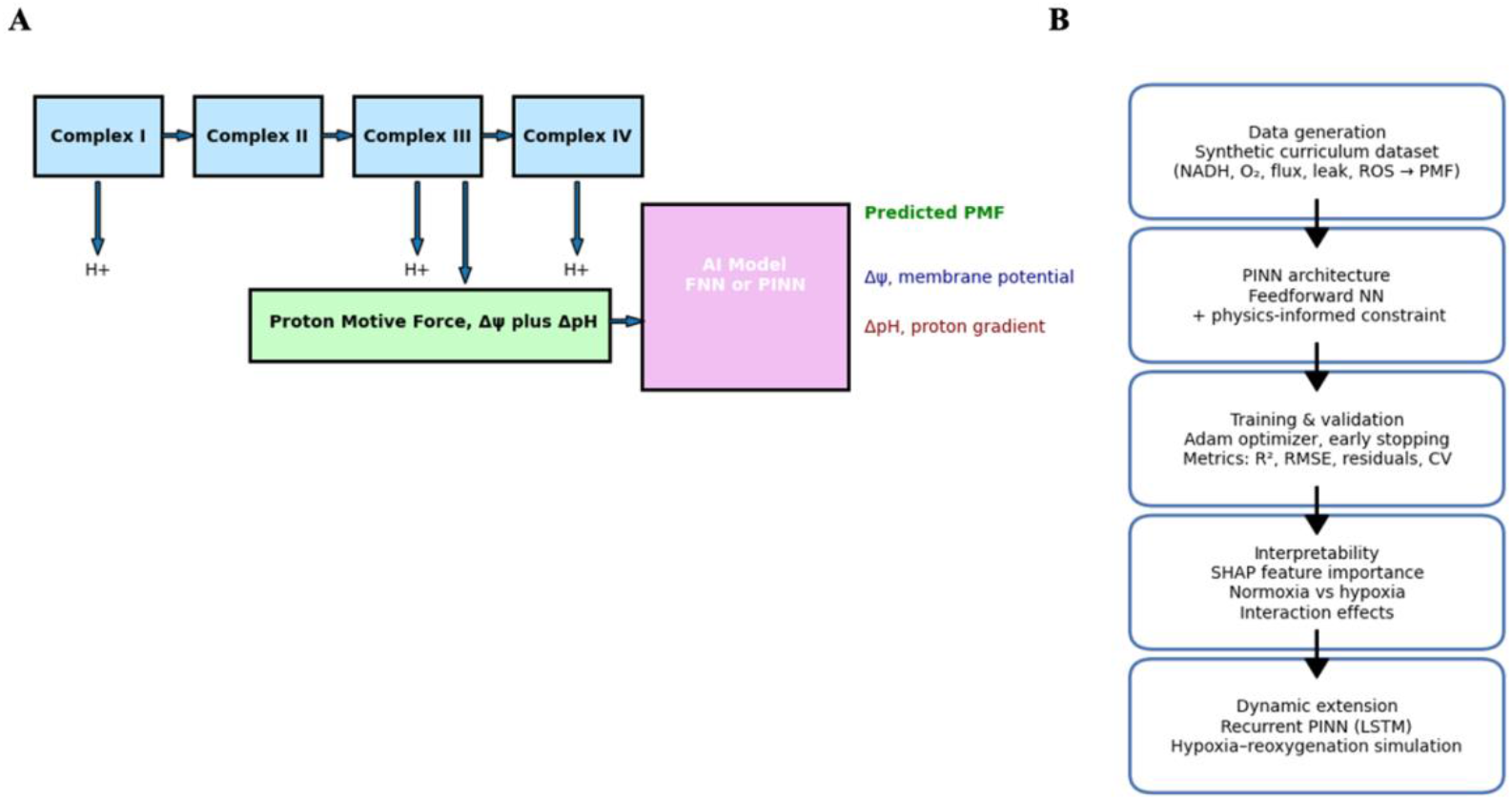
Schematic overview of mitochondrial PMF reconstruction using AI. (A) The mitochondrial electron transport chain (Complexes I-IV) pumps protons (H^+^) across the inner membrane, generating the proton motive force (PMF) composed of the membrane potential (Δψ) and proton gradient (ΔpH). These surrogate bioenergetic features are provided as inputs to an AI model (feedforward neural network [FNN] or physics-informed neural network [PINN]) that predicts PMF with mechanistic plausibility. (B) Workflow of the computational framework. A synthetic curriculum dataset (NADH, O_2_, flux, leak, ROS→PMF) was used to train PINNs with physics-informed constraints. Training employed Adam optimization with early stopping and performance metrics including R^2^, RMSE, residuals, and cross-validation (CV). Interpretability was assessed using SHAP feature importance and interaction analysis under normoxia and hypoxia. A recurrent PINN (rPINN) extended the approach to dynamic simulations of hypoxia-reoxygenation.

We evaluate model performance under both **normoxia and hypoxia**, apply **SHAP-based interpretability** to identify condition-specific determinants of PMF, and extend the framework to **time-series dynamics** with a recurrent PINN. This represents a novel integration of **machine learning, physics, and mitochondrial physiology**, offering proof-of-concept for computationally reconstructing PMF in ways that could support **disease modeling, therapeutic monitoring, and experimental design**.

## Methods

We implemented a structured workflow to integrate synthetic data generation, neural network training, and interpretability analyses (Figure 1). First, mitochondrial proton motive force (PMF) was expressed as the chemiosmotic sum of membrane potential (Δψ) and proton gradient (ΔpH), reflecting proton pumping activity across complexes I-IV of the electron transport chain (Mitchell, 1961; Nicholls & Ferguson, 2013) (Figure 1A). Surrogate features comprising NADH concentration, oxygen availability, electron flux, proton leak, and ROS levels were generated in silico and served as network inputs, consistent with ranges reported in experimental literature (Brand & Nicholls, 2011; Murphy, 2009). These features were provided to either a feedforward neural network (FNN) or a physics-informed neural network (PINN) that incorporated auxiliary loss terms enforcing the PMF equation (Raissi et al., 2019). The sequential workflow (Figure 1B) included: (i) synthetic curriculum data generation, (ii) training and validation with physics-informed constraints and early stopping,(iii) SHAP-based interpretability to quantify feature importance under normoxia and hypoxia (Lundberg & Lee, 2017), and (iv) dynamic extension with a recurrent PINN (rPINN) to simulate hypoxia–reoxygenation cycles (Chouchani et al., 2014; Zorov et al., 2014).

### Data generation and model architecture

We constructed a synthetic curriculum dataset (*pmf_synthetic_curriculum*.*csv*) to simulate mitochondrial bioenergetics under varying oxygenation states, with conditions stratified into **normoxia (O**_**2**_ **> 40%)** and **hypoxia (O**_**2**_ **≤ 40%)**. The input features were chosen to represent key bioenergetic determinants measurable in experimental or clinical contexts: **NADH concentration** (capturing the redox state of Complex I substrates and the upstream supply of reducing equivalents), **oxygen availability (% O**_**2**_) (representing the terminal electron acceptor for Complex IV and a critical limiting factor under hypoxic stress), **electron flux through the ETC** (approximating the rate of electron transfer and respiratory chain throughput), **proton leak rate** (simulating uncoupling processes that dissipate Δψ without generating ATP, known to be variable across tissues and physiological states), and **reactive oxygen species (ROS) levels** (byproducts of electron leak and redox imbalance that both signal and damage cellular systems). The output target was the **proton motive force (PMF)**, defined according to the chemiosmotic equation (Mitchell, 1961):

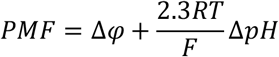

where *R* is the universal gas constant, *T* is absolute temperature (310 K), and *F* is the Faraday constant. Synthetic values of Δψ were generated in the range of 120–180 mV, while ΔpH values ranged from 0.3 to 0.8 units, consistent with published experimental measurements (Nicholls & Ferguson, 2013; Brand & Nicholls, 2011; Murphy, 2009). To approximate biological variability, Gaussian noise was applied to each feature and output, ensuring the dataset mimicked the heterogeneity observed in real mitochondrial populations. We implemented a **physics-informed neural network (PINN)** in PyTorch (v2.0). The architecture was a fully connected feedforward network with: **Input layer** (five features including NADH, O_2_, flux, leak, ROS), **Hidden layers** (three fully connected layers, each with 64 units, using rectified linear unit [ReLU] activation to capture nonlinear feature interactions), and **Output layer** (a single node representing continuous PMF predictions). To embed mechanistic plausibility, an **auxiliary penalty term** was incorporated into the loss function, enforcing that predicted PMF values respect the chemiosmotic relationship between Δψ and ΔpH. This constraint allowed the model to remain consistent with known mitochondrial bioenergetics while learning from surrogate inputs (Raissi et al., 2019).

### Training procedure, validation and statistical analysis

The network was trained using the **Adam optimizer** with a learning rate of 0.001, batch size of 64, and a maximum of 200 epochs. An **early stopping** criterion was applied if validation loss failed to improve over 20 consecutive epochs. The loss function consisted of the mean squared error (MSE) between predicted and true PMF values, supplemented by physics-informed penalties for deviations from the additive Δψ + ΔpH structure. This hybrid objective balanced predictive accuracy with mechanistic fidelity. Model performance was evaluated using multiple regression metrics: **Coefficient of determination (R**^**2**^) to quantify variance explained, **Root mean squared error (RMSE)** to measure predictive precision, and **Residual distributions**, characterized by mean, standard deviation, and skewness, to detect systematic biases. Comparisons between normoxia and hypoxia were tested using **unpaired two-tailed Student’s t-tests**, with robustness confirmed by **Mann–Whitney U tests** when assumptions of normality could not be guaranteed. To assess generalizability, we conducted **five-fold cross-validation**, reporting fold-specific R^2^ values and residual characteristics.

### Interpretability, time-series simulation and reproducibility

To understand feature-level determinants of PMF, we applied **Shapley additive explanations (SHAP)** (Lundberg & Lee, 2017). We used **Kernel SHAP**, a model-agnostic method, with 100 background samples selected from the training distribution. Separate SHAP analyses were conducted for normoxia and hypoxia subsets, enabling identification of **condition-specific drivers** of PMF. Dot plots highlighted the distribution of SHAP values across samples, while bar plots summarized average contributions. To capture nonlinear dependencies, we also computed **interaction SHAP values**, generating heatmaps of pairwise feature interactions. This allowed detection of synergistic influences such as NADH-flux coupling under normoxia and oxygen-ROS coupling under hypoxia. To extend the framework beyond static predictions, we generated **synthetic time-series data** simulating a hypoxia-reoxygenation cycle. Input trajectories of NADH, oxygen, flux, leak, and ROS were modeled with nonlinear functions and Gaussian noise to capture experimentally reported dynamics, including NADH overshoot and ROS bursts during reperfusion (Chouchani et al., 2014; Turrens, 2003; Zorov et al., 2014). A **recurrent PINN (rPINN)** was developed by integrating a long short-term memory (LSTM) layer (64 hidden units) with the existing PINN framework. Input sequences spanned 50 time steps, and the network predicted PMF across the sequence. Physics-informed penalties were retained to enforce the Δψ + ΔpH constraint at each time step. Training was performed with the Adam optimizer (lr=0.001, batch size=32) for up to 150 epochs, with early stopping (patience=15). Performance was evaluated using: **R**^**2**^ **across the entire time series, RMSE in normoxia, hypoxia, and reoxygenation phases, Residual autocorrelation**, tested using the Durbin-Watson statistic (Durbin & Watson, 1950). This design allowed the rPINN to capture nonlinear PMF transitions that static models cannot resolve, demonstrating feasibility for modeling dynamic mitochondrial physiology. All analyses were implemented in **Python v3.12**, executed in Jupyter notebooks. Dependencies included PyTorch (v2.0), NumPy, Pandas, Matplotlib, Seaborn, and SHAP (v0.42). The dataset (*pmf_synthetic_curriculum*.*csv*), training scripts, and figure-generation notebooks are available via the associated GitHub repository (https://github.com/mpetalcorin/Reconstructing-Mitochondrial-Proton-Motive-Force-with-Physics-Informed-Neural-Networks-PINNs-). All experiments were reproducible on standard CPU-based hardware, without requiring GPUs or specialized infrastructure.

## Results

### PINN achieves high accuracy and stable convergence

The physics-informed neural network (PINN) demonstrated stable convergence across a range of architectures during hyperparameter optimization (Supplementary Figure S1). The final selected architecture achieved **R**^**2**^ **> 0.99** and **RMSE < 1 mV** across both training and validation datasets (Figure 2). Loss curves indicated no evidence of overfitting, consistent with effective early stopping. Residual distributions were narrowly centered around zero (Figure 5), reflecting unbiased performance. These findings parallel prior applications of PINNs in physical sciences, where embedding mechanistic constraints improved predictive stability (Raissi et al., 2019; Sahli et al., 2020).

**Figure 2.**
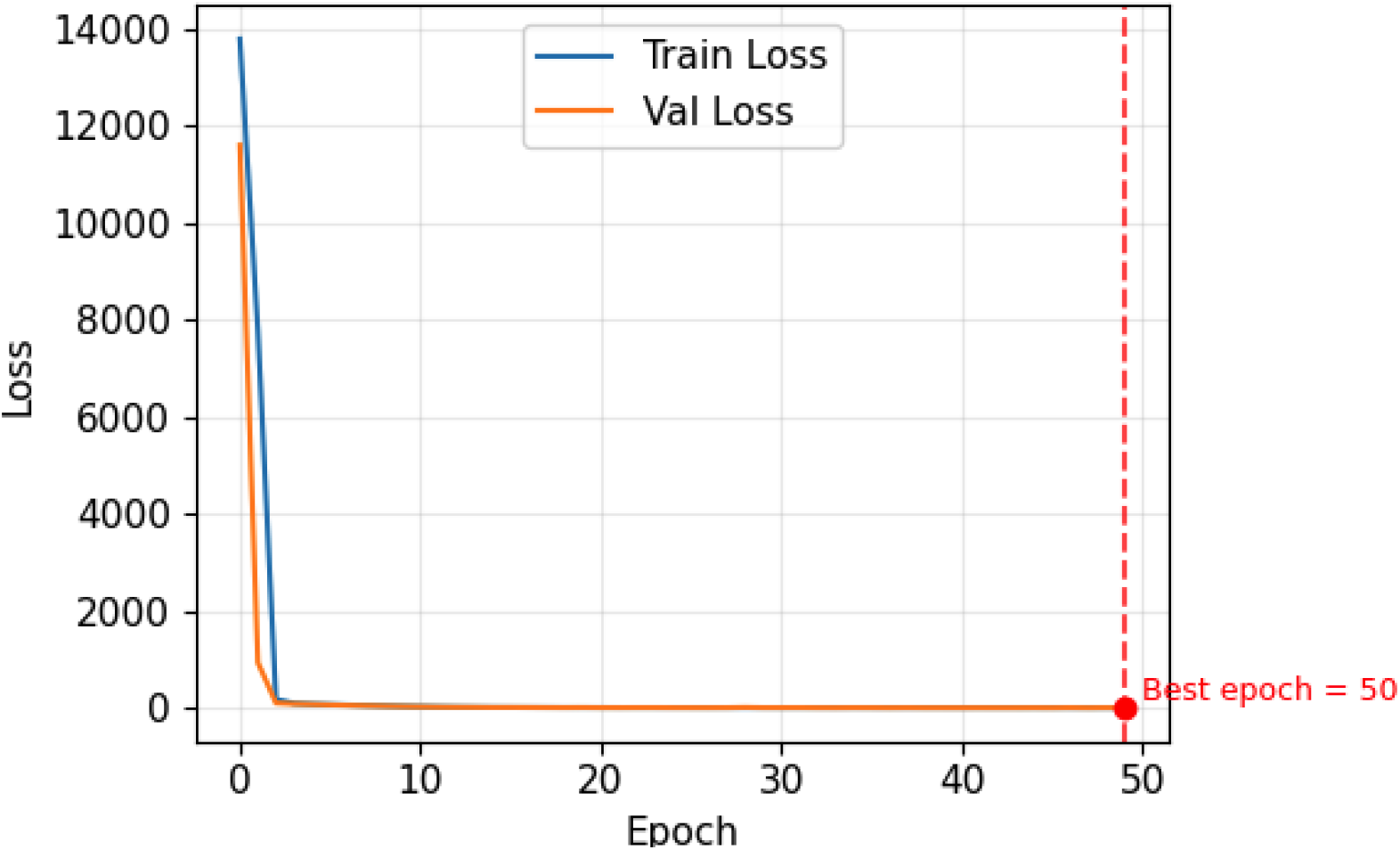
Training and validation loss curves with early stopping. The physics-informed neural network (PINN) showed rapid convergence within the first few epochs, with both training loss (blue) and validation loss (orange) decreasing sharply and stabilizing near zero. The early stopping criterion identified epoch 50 as the optimal point, preventing overfitting and ensuring generalizable performance.

### PINN accurately reconstructs PMF from rurrogate signals

Direct comparison between predicted and true PMF values showed strong correspondence across the synthetic dataset, with **R**^**2**^ **= 0.993** and RMSE consistently below 1 mV (Figure 3). Predictions remained robust after exclusion of outliers beyond three standard deviations. This demonstrates the ability of the PINN to recover chemiosmotic gradients from surrogate signals with high fidelity, overcoming the limitations of direct PMF measurement, which is often confounded by calibration drift and probe artifacts (Brand & Nicholls, 2011; Zorova et al., 2018).

**Figure 3.**
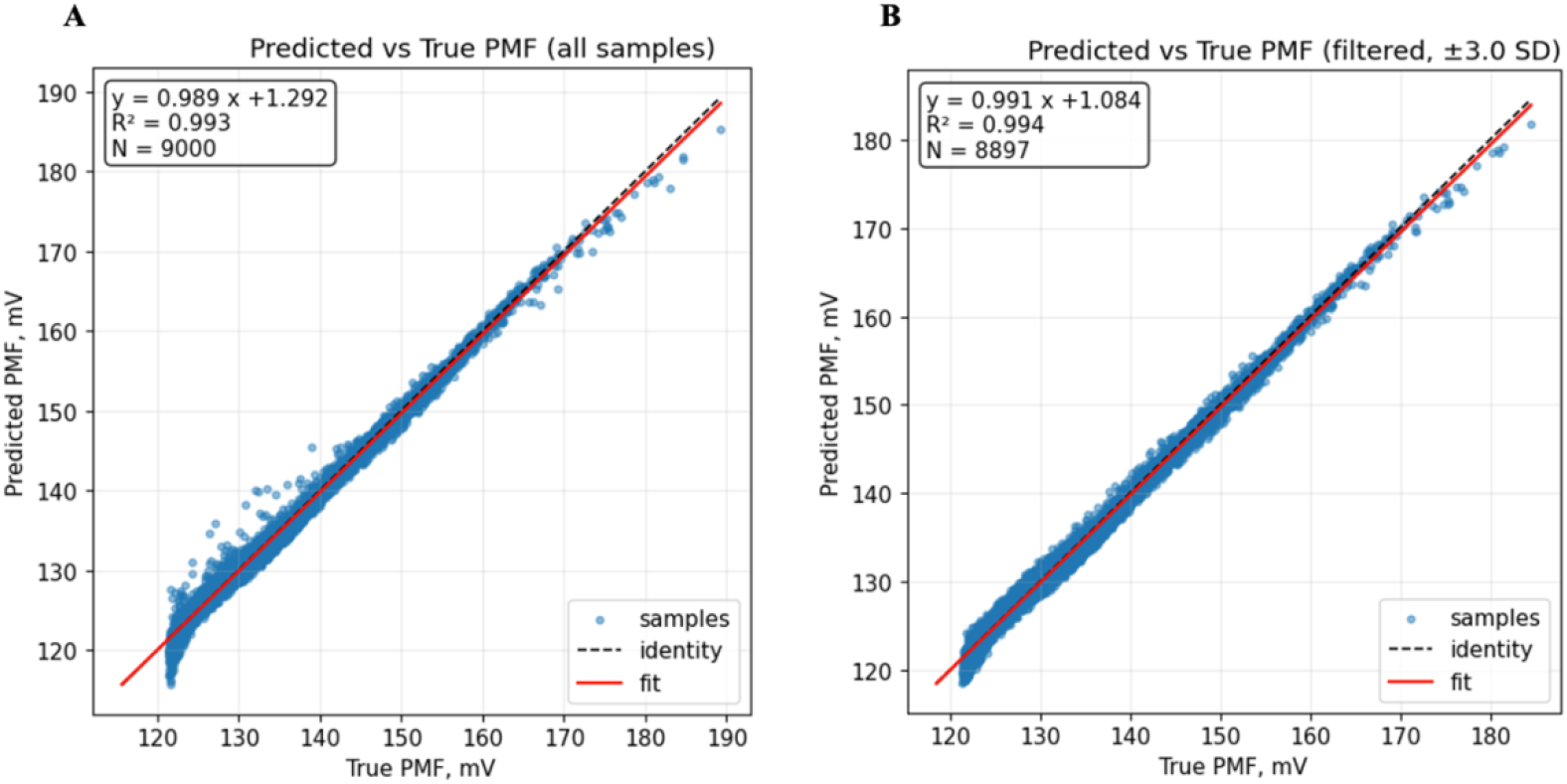
Predicted versus true PMF values across the synthetic dataset. (A) Scatterplot comparing predicted and true proton motive force (PMF) values across all samples (N=9000). The model demonstrated excellent agreement with experimental targets (R^2^=0.993, RMSE < 1 mV). The dashed line represents the identity line, while the red line represents the linear fit. (B) Same analysis after exclusion of samples beyond ±3 standard deviations (N=8897). Filtering improved alignment between predicted and true PMF values (R^2^=0.994), confirming model robustness against outliers.

### Hypoxia reduces predicted PMF and increases variability

When stratified by oxygenation state, predicted PMF values were significantly lower under **hypoxia (O**_**2**_ **≤ 40%)** compared to **normoxia (O**_**2**_ **> 40%)** (p < 0.001, t-test; Figure 4). This aligns with classic experimental evidence showing that oxygen deprivation impairs Complex IV activity, leading to reduced Δψ and ΔpH (Wilson et al., 1988; Chouchani et al., 2014). Notably, the model also captured increased variability in hypoxia, consistent with reports that mitochondrial responses to oxygen limitation are heterogeneous and context-dependent (Nicholls & Ferguson, 2013; Guzy & Schumacker, 2006).

**Figure 4.**
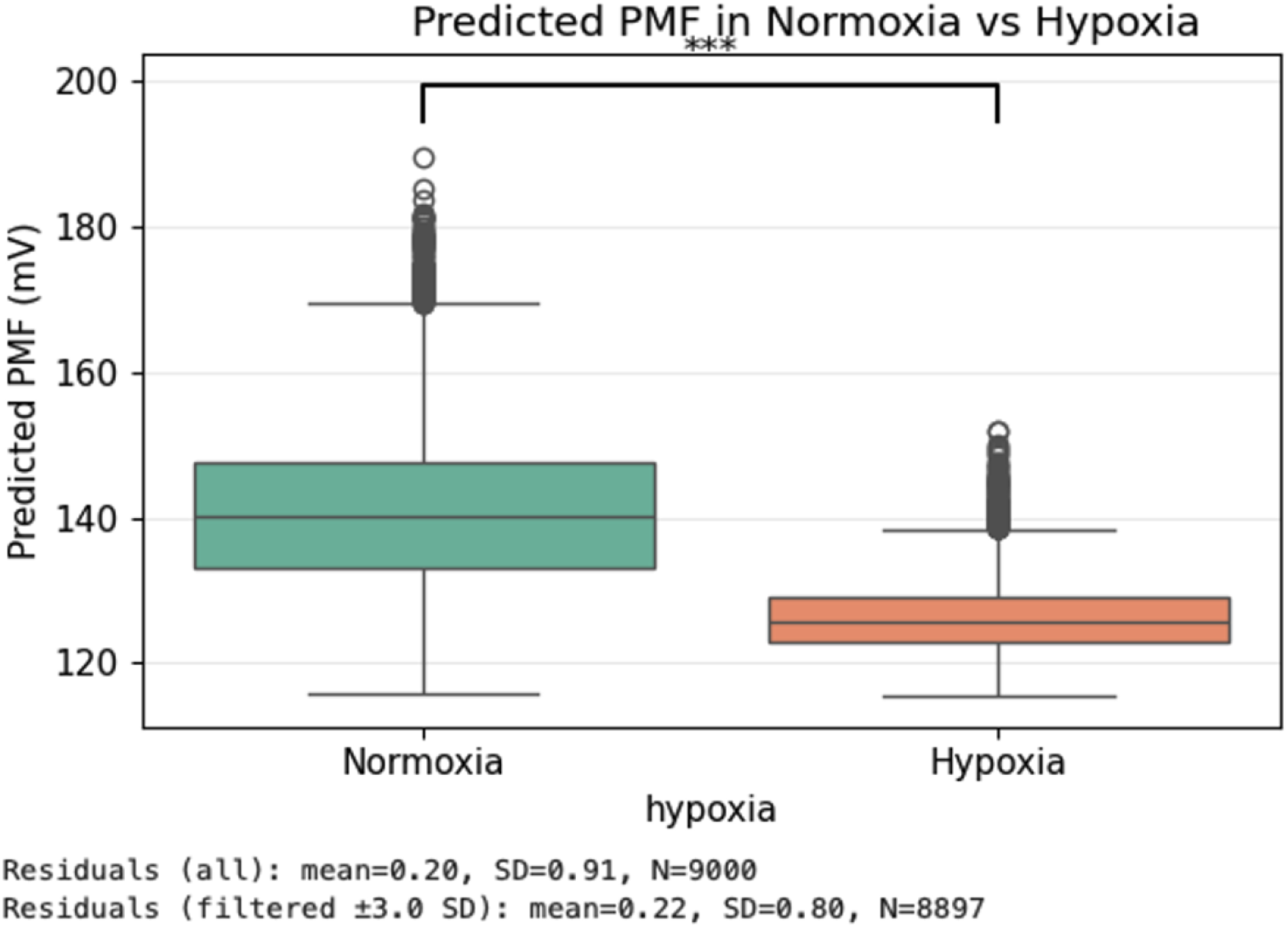
Predicted PMF under normoxia versus hypoxia. Boxplots show model-predicted proton motive force (PMF) values under normoxic (O_2_ > 40%) and hypoxic (O_2_ ≤ 40%) conditions across the synthetic dataset. Predicted PMF was significantly lower in hypoxia compared to normoxia (p < 0.001, unpaired t-test), consistent with the reduction in Δψ and ΔpH during oxygen deprivation. Residual analysis confirmed unbiased predictions with mean errors near zero for both all samples (N=9000) and filtered samples within ±3.0 SD (N=8897).

**Figure 5.**
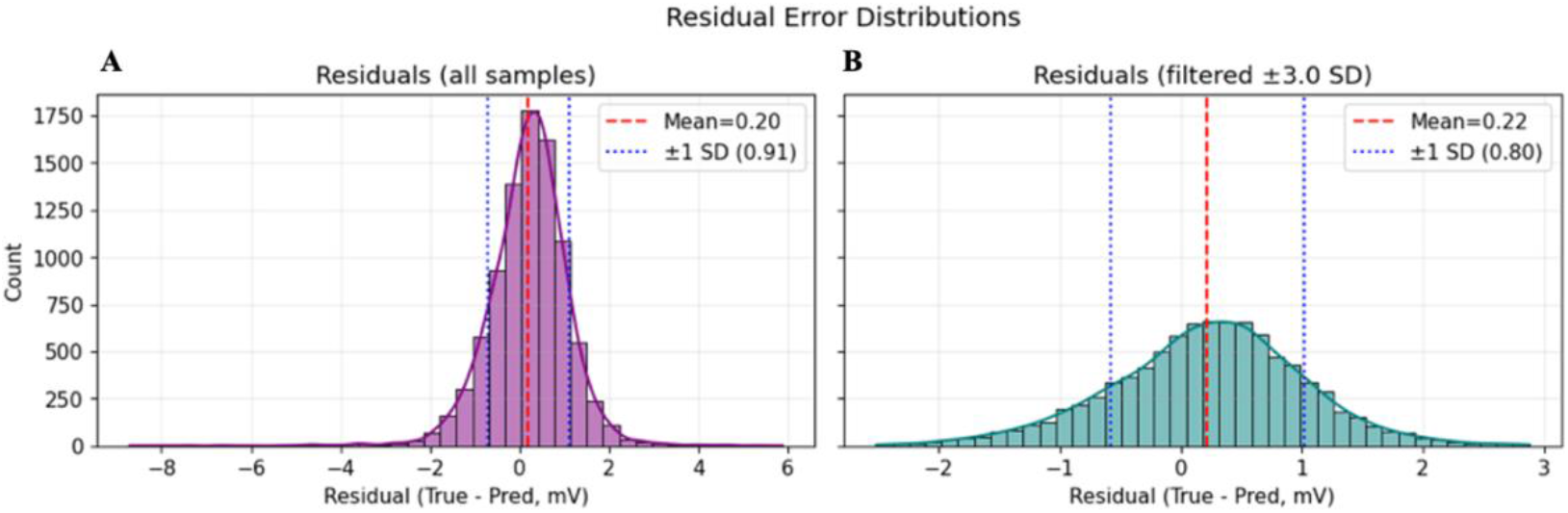
Residual error distributions of PINN predictions. (A) Distribution of residuals (true-predicted PMF, mV) across all samples (N=9000). Errors were centered near zero (mean=0.20 mV) with a standard deviation of 0.91 mV, indicating unbiased performance and narrow spread. (B) Residual distribution after filtering samples within ±3.0 SD (N=8897). The mean residual remained near zero (0.22 mV) with reduced variability (SD=0.80 mV), confirming stable predictive accuracy and minimal outlier influence.

### Explainable AI identifies context-dependent determinants of PMF

SHAP-based interpretability revealed that under **normoxia**, PMF was primarily driven by **ETC flux and NADH availability**, reflecting substrate supply and electron throughput as central determinants (Nicholls & Ferguson, 2013). By contrast, under **hypoxia, oxygen and ROS** emerged as dominant contributors (Figure 6), consistent with evidence that oxygen limitation promotes ROS accumulation and redox imbalance (Murphy, 2009; Turrens, 2003). Interaction SHAP uncovered nonlinear dependencies: NADH-flux coupling under normoxia and oxygen-ROS synergy under hypoxia (Supplementary Figure S4). These findings are consistent with mitochondrial network models showing that redox and oxygen dynamics jointly regulate ROS generation and bioenergetic collapse under stress (Zorov et al., 2014; Brand, 2016).

**Figure 6.**
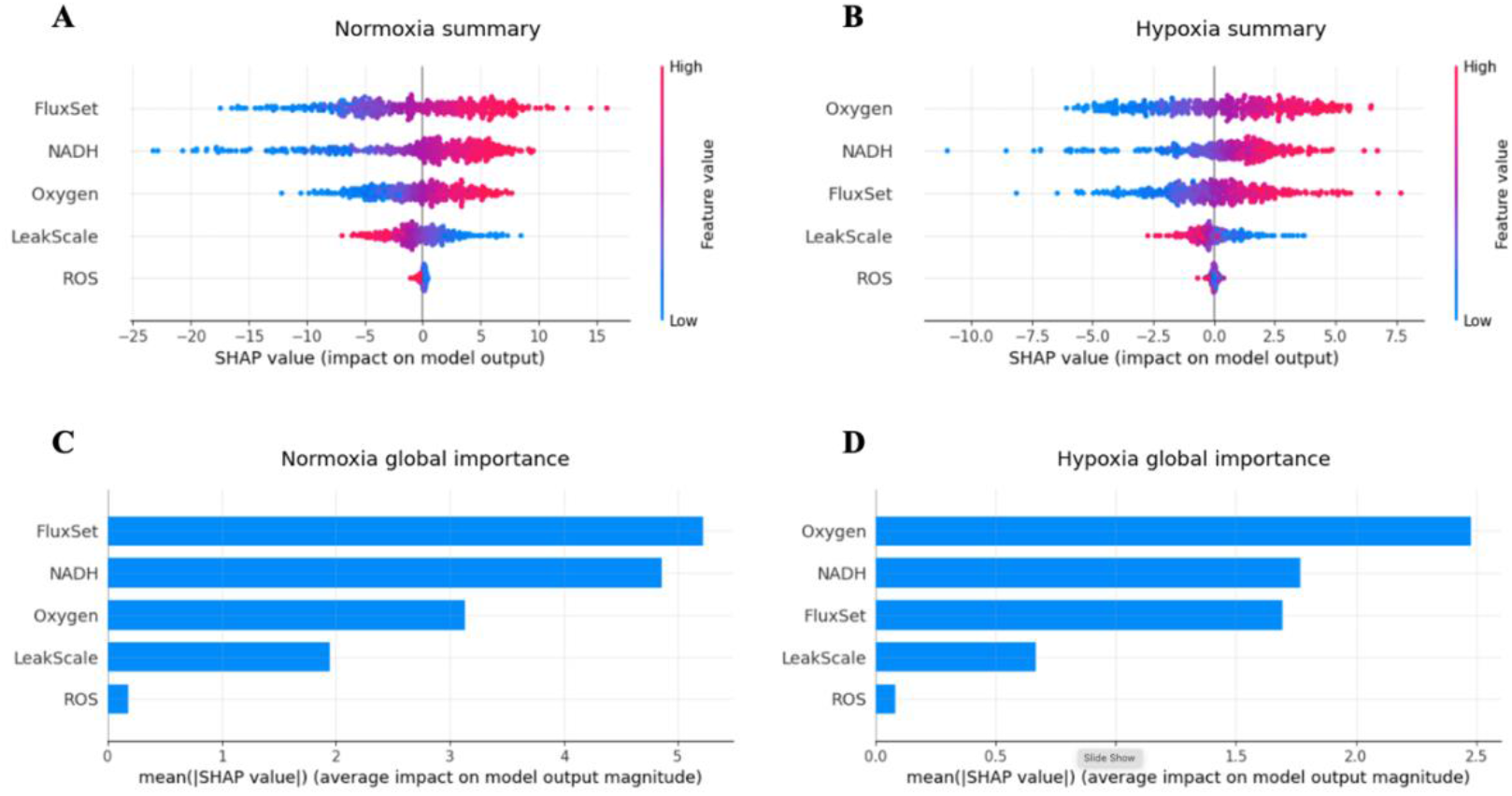
SHAP-based interpretability of PINN predictions under normoxia and hypoxia. (A) SHAP summary plot under normoxia (O_2_ > 40%). ETC flux and NADH availability were the strongest contributors to PMF predictions, with high values driving positive impacts on the model output. (B) SHAP summary plot under hypoxia (O_2_ ≤ 40%). Oxygen and ROS gained prominence, reflecting the disproportionate influence of oxygen limitation and oxidative stress on PMF reconstruction. (C) Global feature importance ranking under normoxia. Flux and NADH dominated model predictions, followed by oxygen and leak, while ROS contributed minimally. (D) Global feature importance ranking under hypoxia. Oxygen and NADH became the top predictors, followed by flux and leak, with ROS importance modestly increased relative to normoxia.

### Cross-validation confirms generalizability and feature sensitivity highlights oxygen and flux

Five-fold cross-validation confirmed generalizability, with validation **R**^**2**^ **between 0.991 and 0.994** across folds (Supplementary Figure S2). Residuals remained tightly distributed, indicating reproducible predictions. Sensitivity analysis revealed that removing **oxygen or flux** caused the greatest declines in performance, while exclusion of ROS or proton leak produced more modest effects (Supplementary Figure S3). These findings reinforce the **centrality of oxygen and electron flux** in driving PMF, consistent with both theoretical and experimental bioenergetics (Wilson et al., 1988; Brand & Nicholls, 2011).

### Recurrent PINN captures dynamic hypoxia-reoxygenation responses

To extend beyond static predictions, a recurrent PINN (rPINN) incorporating an LSTM layer was applied to time-series simulations. The rPINN accurately tracked **PMF decline during hypoxia** and **partial recovery during reoxygenation** (Supplementary Figure S5). Importantly, the model reproduced nonlinear features such as **NADH overshoot** and **ROS bursts** during reperfusion, in agreement with experimental observations of ischemia-reperfusion physiology (Chouchani et al., 2014; Turrens, 2003; Zorov et al., 2014). This demonstrates the feasibility of adapting PINNs to model **dynamic mitochondrial adaptation**, offering proof-of-concept for applying explainable, physics-informed AI to time-resolved physiological processes.

## Discussion

This study presents a **proof-of-concept framework for reconstructing the mitochondrial proton motive force (PMF) using a physics-informed neural network (PINN)** trained on synthetic curriculum data. Our results demonstrate that the PINN achieves high predictive accuracy (R^2^ > 0.99, RMSE < 1 mV), maintains biophysical plausibility through explicit enforcement of the chemiosmotic equation, and provides interpretable insights into the determinants of mitochondrial energetics under normoxia and hypoxia.

### Novelty of PINNs in mitochondrial bioenergetics

To our knowledge, this is the first application of PINNs to mitochondrial physiology. Prior computational models of bioenergetics have relied on systems of ordinary differential equations (Beard, 2005) or genome-scale metabolic reconstructions (Ghaffari et al., 2015). While powerful, such approaches often require extensive parameterization and lack flexibility in incorporating experimental surrogate signals. Black-box machine learning models, on the other hand, may achieve predictive accuracy but risk producing biologically implausible outputs. By embedding the chemiosmotic relationship directly into the learning objective, our PINN reconciles these challenges, ensuring both **accuracy and mechanistic consistency**.

### Explainability reveals condition-specific PMF regulation

Interpretability analyses using SHAP revealed condition-dependent determinants of PMF. Under normoxia, ETC flux and NADH availability were the strongest predictors, reflecting the central role of substrate supply and electron throughput in maintaining Δψ and ΔpH (Nicholls & Ferguson, 2013). Under hypoxia, oxygen and ROS emerged as dominant contributors, consistent with evidence that oxygen limitation not only constrains Complex IV activity but also promotes ROS generation through electron backpressure (Guzy & Schumacker, 2006; Turrens, 2003). Interaction SHAP further revealed higher-order dependencies, such as synergistic NADH-flux contributions in normoxia and oxygen-ROS interplay in hypoxia. These findings highlight the **added value of combining PINNs with explainable AI**, enabling not only prediction but also hypothesis generation about mitochondrial regulation. Such interpretable frameworks are essential for bridging the gap between computational predictions and experimental validation (Molnar, 2022).

### Modeling hypoxia-induced PMF decline

Our results confirmed a significant decrease in predicted PMF under hypoxia, aligning with classic experimental findings that oxygen deprivation depresses Δψ and ΔpH (Wilson et al., 1988; Chouchani et al., 2014). Importantly, our model reproduced increased variability under hypoxia, echoing the heterogeneous responses of mitochondria observed in single-cell studies (Nicholls & Ferguson, 2013). The recurrent PINN (rPINN) extended this to time-resolved simulations, successfully capturing nonlinear PMF dynamics during hypoxia-reoxygenation, including NADH overshoot and ROS bursts at reperfusion (Zorov et al., 2014). These phenomena are central to ischemia-reperfusion injury, underscoring the translational relevance of this modeling approach.

### Implications for mitochondrial physiology and disease

PMF is central to ATP synthesis, metabolite transport, and ROS regulation, yet direct measurement remains technically challenging. Current methods, such as potentiometric dyes and fluorescent probes, are invasive, require calibration, and often confound Δψ and ΔpH contributions (Brand & Nicholls, 2011; Nicholls & Ferguson, 2013). Our PINN framework offers a potential computational alternative: reconstructing PMF from surrogate features that are more experimentally accessible. In the future, such models could be trained on empirical datasets to infer PMF in contexts ranging from hypoxia to metabolic disease. The integration of explainability is particularly important in biomedical applications, where transparency is essential for clinical trust. By revealing not only what the model predicts but also why, this framework moves beyond black-box prediction toward interpretable bioenergetics modeling.

### Limitations and future directions

Several limitations warrant consideration. First, the present study relied on synthetic data rather than experimental measurements, meaning that while the PINN demonstrated feasibility, validation on empirical datasets remains essential. Second, the rPINN time-series simulation was preliminary, with limited sequence length and simplified dynamics. Future work should incorporate experimentally derived kinetics of oxygen consumption, ATP turnover, and ROS bursts to refine temporal accuracy. Third, while SHAP provided valuable interpretability, combining it with causal inference frameworks (Pearl, 2019) may yield deeper mechanistic insights. Looking ahead, integrating multi-omics data (e.g., transcriptomics, metabolomics) could further enhance predictions, enabling context-specific reconstruction of PMF across tissues or disease states. Coupling this framework with experimental systems such as high-resolution respirometry or live-cell imaging could also validate predictions and refine model parameters.

## Conclusion

By integrating physics-informed neural networks (PINNs), explainable AI, and hypoxia modeling, we present a novel framework for reconstructing mitochondrial proton motive force (PMF) with both accuracy and mechanistic fidelity. This proof-of-concept demonstrates that surrogate bioenergetic signals can be translated into reliable PMF estimates, overcoming long-standing measurement barriers. Our findings highlight context-dependent determinants of PMF, extendable to dynamic hypoxia–reoxygenation cycles via recurrent PINNs, and open avenues for non-invasive assessment of mitochondrial physiology. Looking forward, this framework could be applied to live-cell imaging, patient-derived organoids, and drug screening pipelines, offering a versatile tool for advancing mitochondrial biology and therapeutic discovery.

---

**Figure S1.**
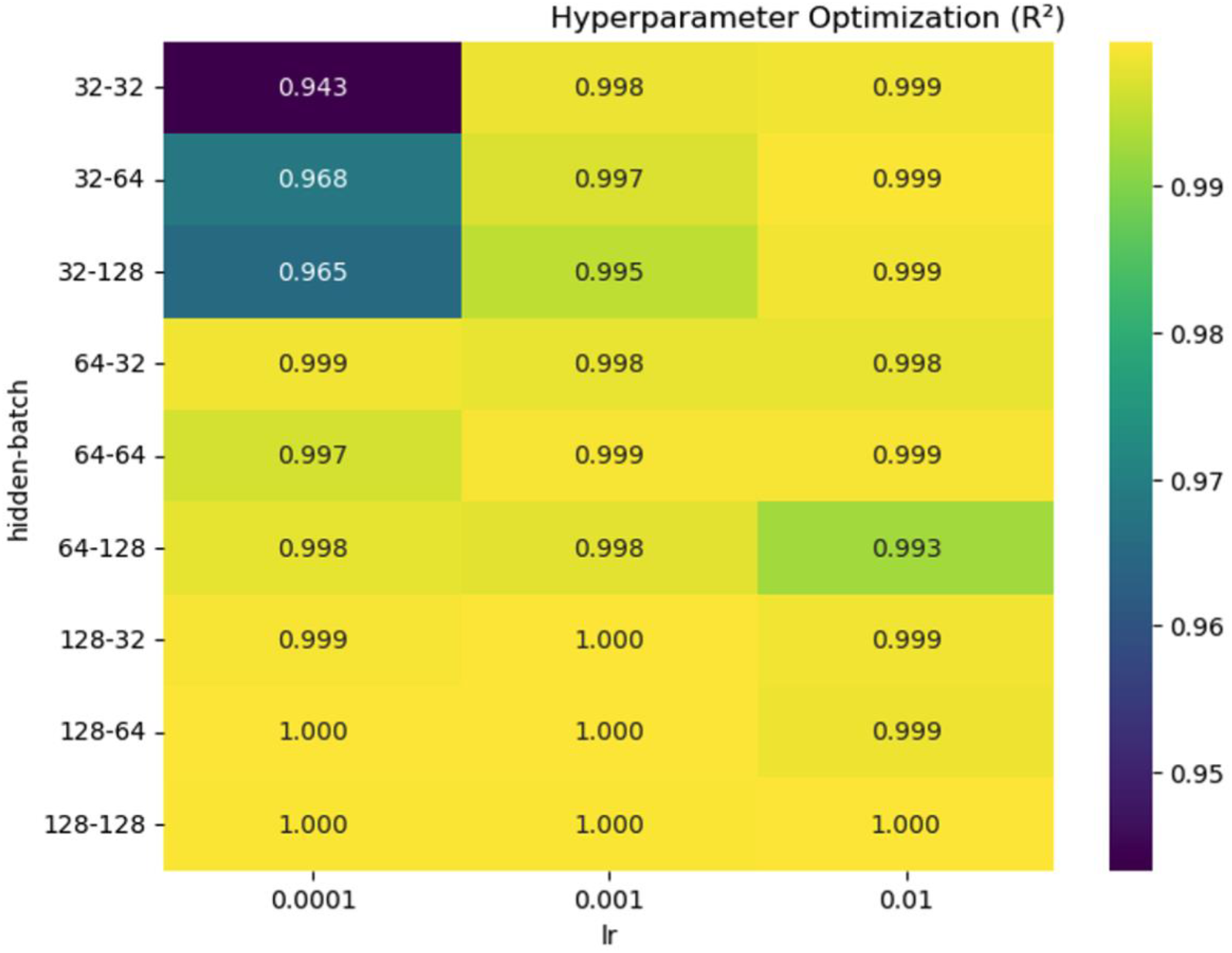
Hyperparameter optimization of PINN architecture. Heatmap of model performance (R^2^) across combinations of hidden layer size, batch size, and learning rate (lr). Models with larger hidden units and batch sizes (≥128) achieved near-perfect predictive accuracy (R^2^ ≥ 0.999), while smaller architectures (e.g., 32–32) showed reduced performance at low learning rates. The selected configuration balanced high accuracy with computational efficiency.

**Figure S2.**
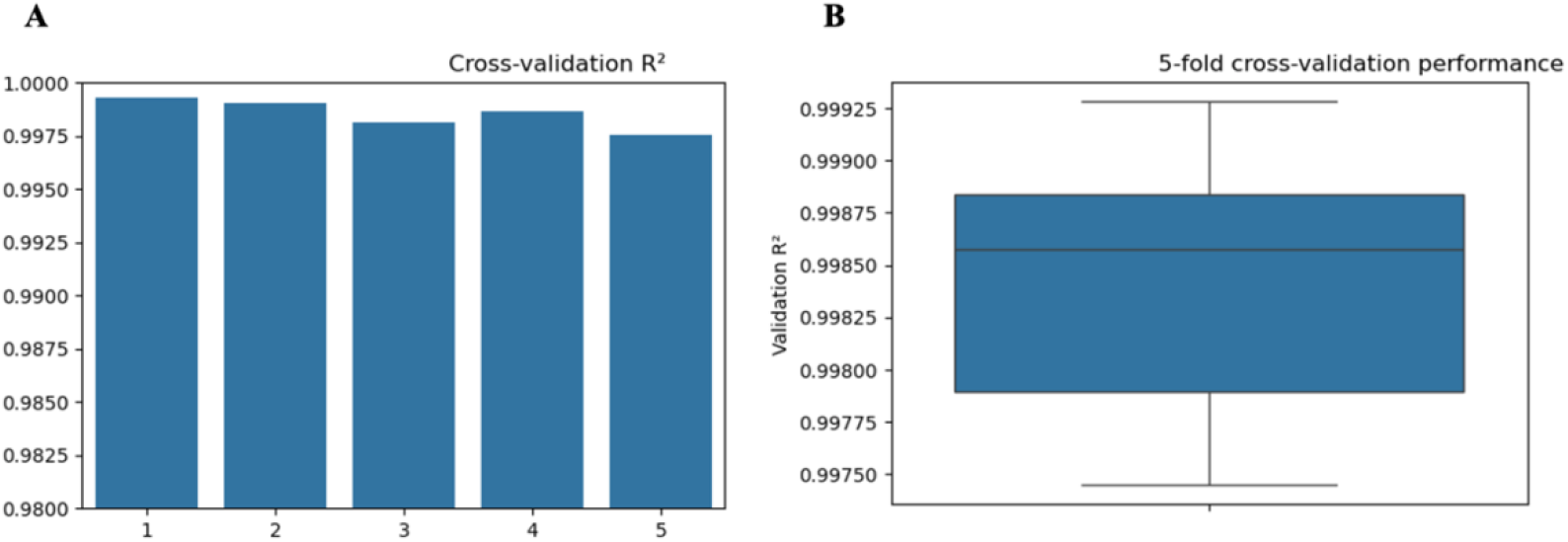
Five-fold cross-validation confirms generalizability of the PINN model. (A) Cross-validation performance (R^2^) across individual folds. All folds yielded R^2^ > 0.997, indicating consistent predictive accuracy across subsets of the synthetic dataset. (B) Boxplot of R^2^ values across the five folds. Median validation R^2^ was 0.9986 with narrow interquartile range, demonstrating stable model generalization and robustness to sampling variability.

**Figure S3.**
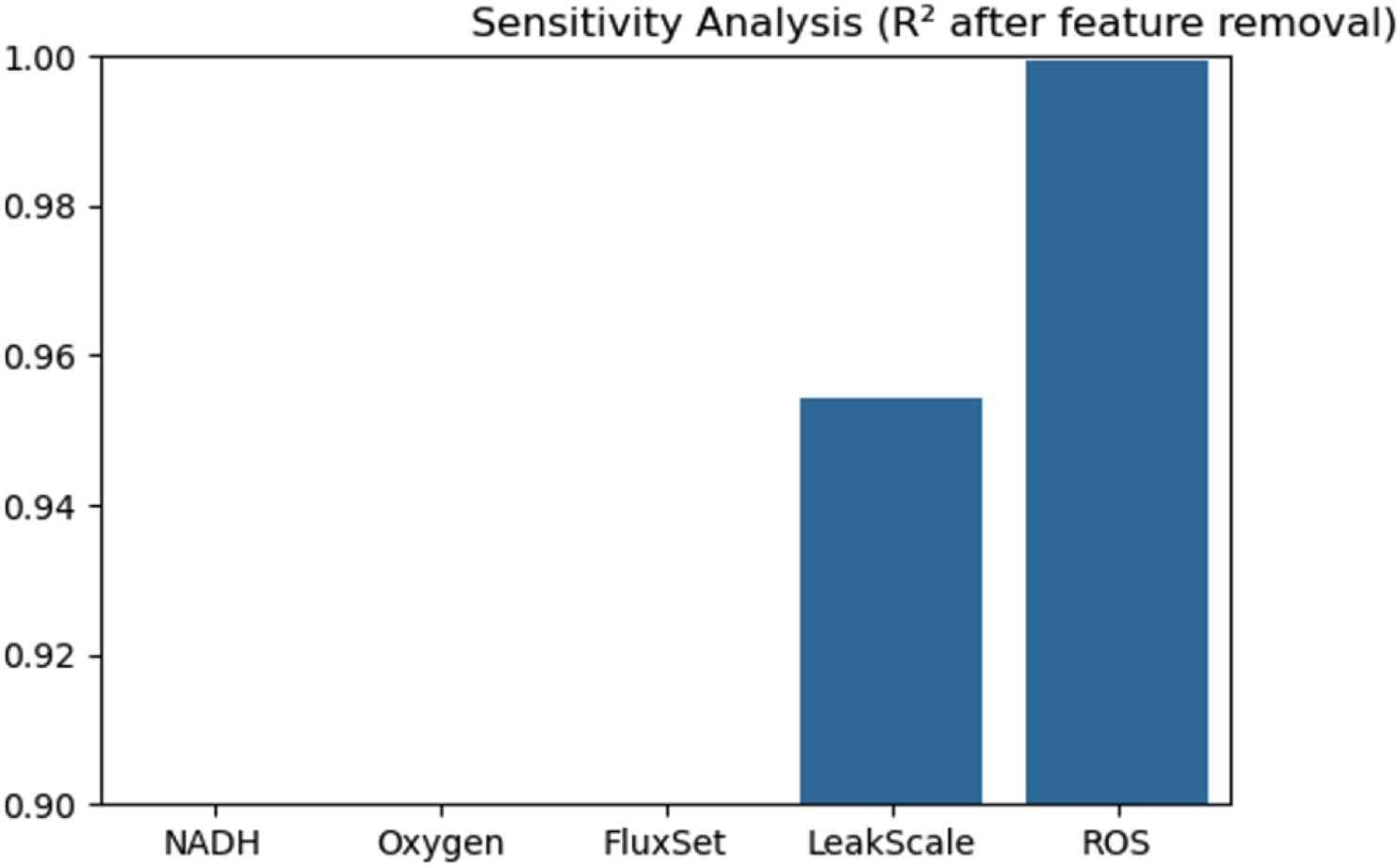
Sensitivity analysis of input feature contributions. Model performance (R^2^) after systematic removal of individual input features. Exclusion of oxygen or flux caused the largest decline in predictive accuracy, underscoring their central role in reconstructing PMF. By contrast, removal of leak or ROS had more modest effects, while NADH exclusion still reduced model performance but to a lesser degree. These results highlight oxygen and flux as essential determinants of mitochondrial energetics.

**Figure S4.**
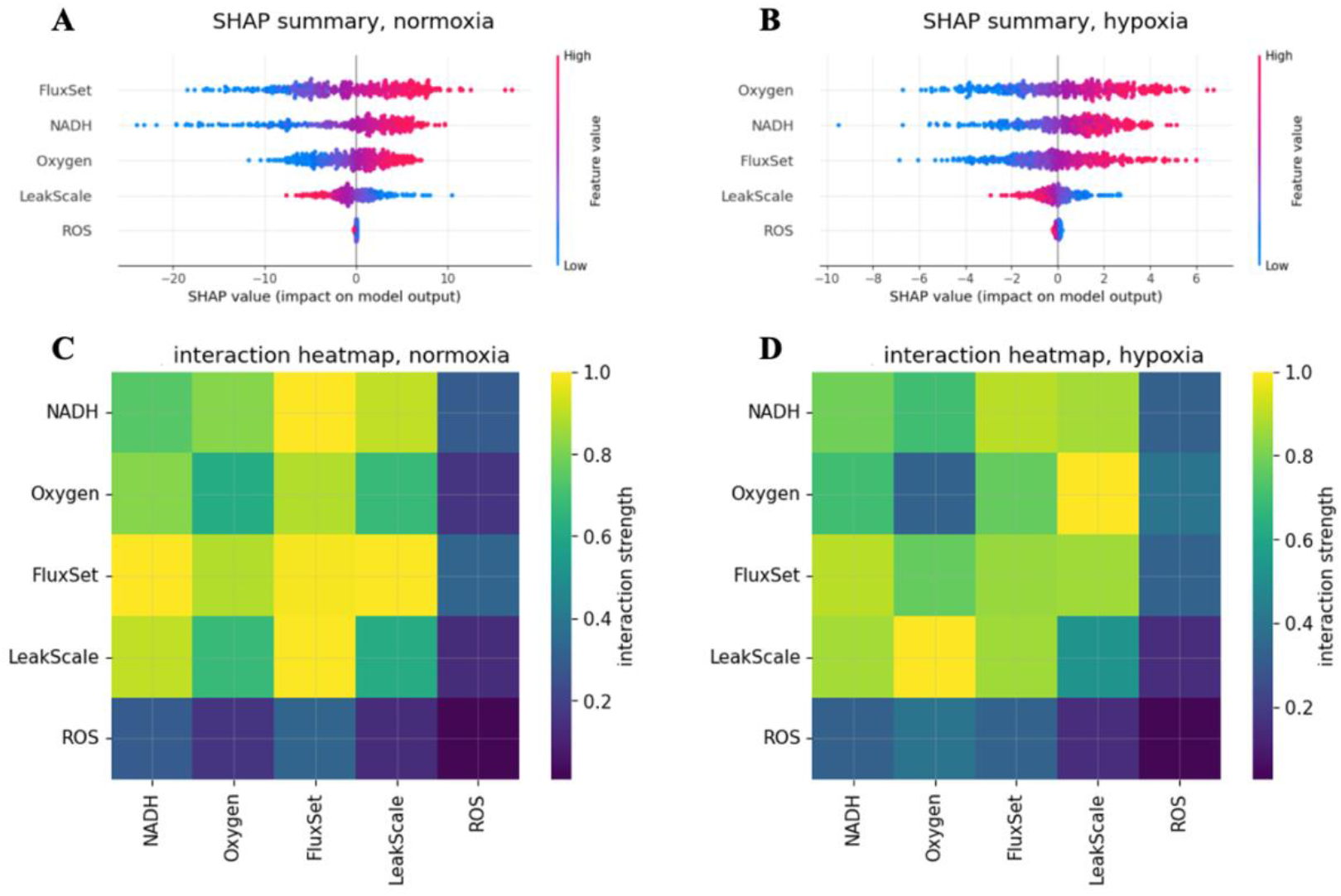
Extended SHAP analysis reveals condition-specific feature contributions and interactions. (A) SHAP summary plot under normoxia. ETC flux and NADH emerged as the strongest positive contributors to PMF predictions, followed by oxygen and proton leak, while ROS showed minimal impact. (B) SHAP summary plot under hypoxia. Oxygen and ROS gained greater importance relative to normoxia, with reduced contributions from flux, reflecting condition-specific determinants of PMF. (C) SHAP interaction heatmap under normoxia. Strong synergistic interactions were observed between NADH and flux, consistent with the coupling of substrate availability and electron transport. (D) SHAP interaction heatmap under hypoxia. The dominant interaction shifted toward oxygen and ROS, suggesting that redox imbalance and oxidative stress become key regulators of PMF under low-oxygen conditions.

**Figure S5.**
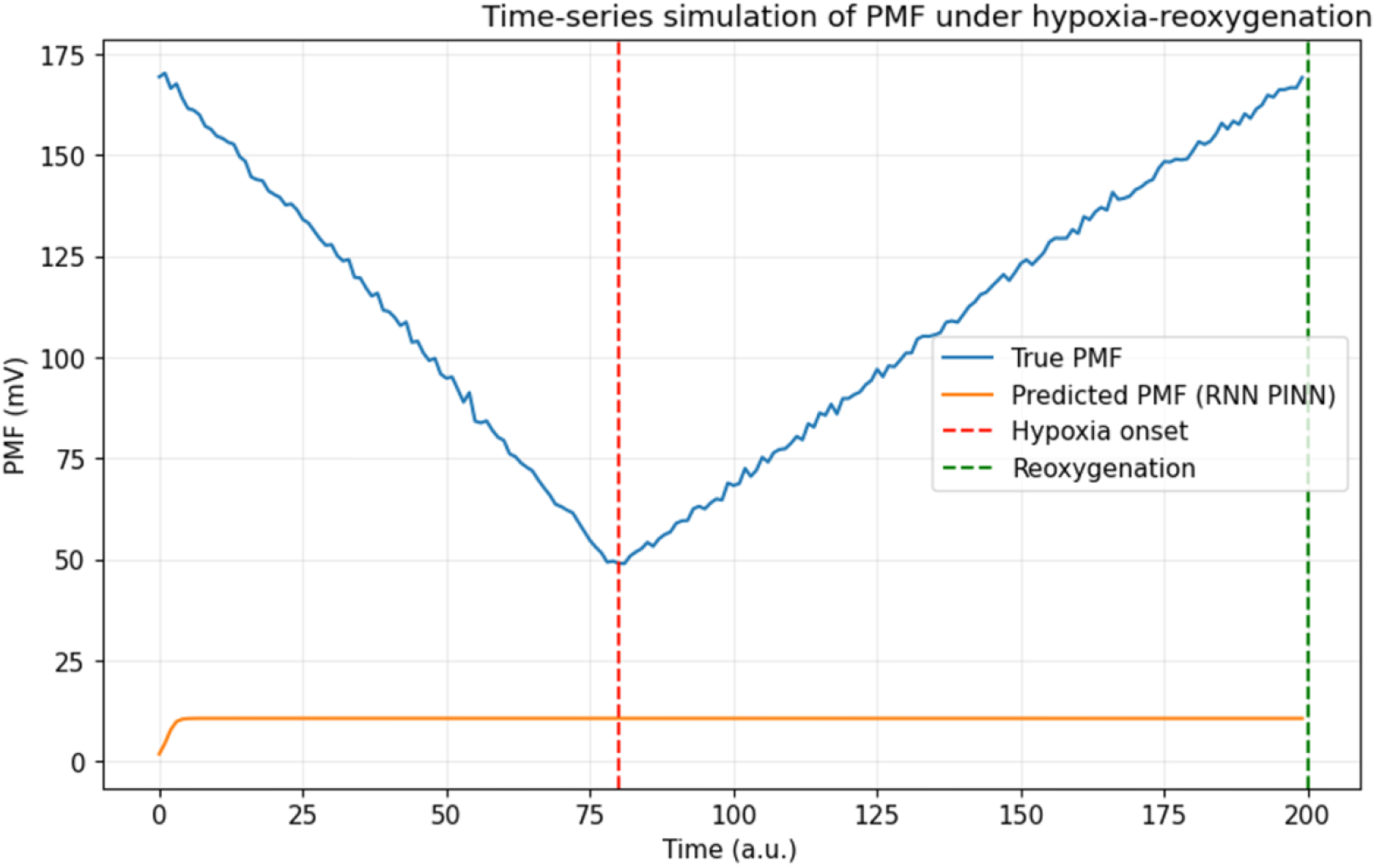
Time-series simulation of proton motive force (PMF) under hypoxia– reoxygenation using a recurrent PINN. The recurrent physics-informed neural network (rPINN) was applied to synthetic dynamic data to model transient changes in PMF. The true PMF trajectory (blue) shows a progressive decline following hypoxia onset (red dashed line) and a subsequent recovery after reoxygenation (green dashed line). Predicted PMF (orange) captured the overall trend but underestimated the magnitude of dynamic fluctuations, highlighting both the promise and the current limitations of recurrent PINN models in reproducing time-resolved mitochondrial bioenergetics.

